# Seipin accumulates and traps diacylglycerols and triglycerides in its ring-like structure

**DOI:** 10.1101/2020.10.27.357079

**Authors:** Valeria Zoni, Wataru Shinoda, Stefano Vanni

## Abstract

Lipid droplets (LD) are intracellular organelles responsible for lipid storage, and they emerge from the endoplasmic reticulum (ER) upon the accumulation of neutral lipids, mostly triglycerides (TG), between the two leaflets of the ER membrane. LD biogenesis takes place at ER sites that are marked by the protein seipin, which subsequently recruits additional proteins to catalyse LD formation. Deletion of seipin, however, does not abolish LD biogenesis, and its precise role in controlling LD assembly remains unclear. Here we use molecular dynamics simulations to investigate the molecular mechanism through which seipin promotes LD formation. We find that seipin clusters TG molecules inside its unconventional ring-like oligomeric structure, and that both its luminal and transmembrane regions contribute to this process. Diacylglycerol, the precursor of TG, also clusters inside the seipin oligomer, in turn promoting TG accumulation. Our results suggest that seipin remodels the membrane of specific ER sites to prime them for LD biogenesis.

**Significance statement:** Metabolic disorders related to aberrant fat accumulation, including lipodystrophy and obesity, are a particularly serious health concern. In cells, fat accumulates in intracellular organelles, named lipid droplets (LDs). LDs form in the endoplasmic reticulum, where triglycerides, the most abundant form of fat, is produced. The Bernardinelli-Seip congenital lipodystrophy type 2 protein, seipin, has been identified as a key regulator of LD formation, but its mechanism of action remains debated and its molecular details mostly obscure. Here, we use molecular dynamics simulations to investigate the mechanism of seipin. We find that seipin can cluster and trap both triglycerides and its precursor, diacylglycerol. Our results suggest that seipin organizes the lipid composition of specific ER sites to prime them for LD biogenesis.

## Introduction

Lipid Droplets (LDs) are the intracellular organelles responsible for fat accumulation(1). As such, they play a central role in lipid and cellular metabolism(1–4) and they are crucially involved in metabolic diseases such as lipodystrophy and obesity(5–7).

Formation of LDs occurs in the endoplasmic reticulum (ER), where neutral lipids (NLs), namely triglycerides (TG) and cholesteryl esters, constituting the core of LDs are synthesized by acyltransferases that are essential for LD formation(8). The current model of LD formation posits that NLs are stored between the two leaflets of the ER bilayer, where they aggregate in nascent oblate lens-like structure with diameters of 40-60 nm(9), before complete maturation and budding towards the cytosol(10–13). Recent experiments suggest that LDs form at specific ER sites marked by the protein seipin(14), upon arrival of its interaction partner protein promethin/LDAF1 (LDO in yeast) (15–19). These recent observations confirm previous work showing that seipin, in addition to modulating LD budding and growth(14, 20–22) and LD-ER contacts(21, 23), is also a major player in the early stages of LD formation, as deletion of seipin leads to TG accumulation in the ER and a delay in the formation of, possibly aberrant, LDs(20, 24).

The role of seipin in LD formation is potentially coupled to its function in regulating lipid metabolism(25–27), and notably that of phosphatidic acid (PA)(28–30). Recently, seipin-positive ER loci have been shown to be part of a larger protein machinery that also includes membrane and lipid remodeling proteins of the TG synthesis pathway(31), most notably, Lipin (Pah1 in yeast) and FIT proteins (Yft2 and Scs3 in yeast), for which PA is either a known substrate (Lipin/Pah1)(32) or a likely one (FIT/Yft2/Scs3)(33).

Despite this thorough characterization of the cellular role of seipin in LD formation, the molecular details of its mechanism remain mostly unclear. Recently, the structure of the luminal part of the seipin oligomer has been solved at 3.7-4.0 Å resolution using electron microscopy(28, 34), paving the way for the investigation of the relationship between its three dimensional structure and its mode of action. These studies revealed that the luminal domain of seipin consists of an eight-stranded beta sandwich, together with a hydrophobic helix (HH) positioned toward the ER bilayer. Notably, the seipin oligomer assembles into a ring-like architecture, an unconventional assembly in lipid bilayers that rather resembles the shape of microbial pore-forming assemblies(35) or GroEL-GroES chaperones(36, 37).

From a stochiometric point of view, both fluorescence and electron microscopy data are consistent with the presence of a single seipin oligomer per nascent LD (14, 15). Hence, the structure of the luminal part of seipin is consistent with two proposed modes of action: seipin could mark the sites of LD formation by controlling TG flow in and out of the nascent droplet(14), or, alternatively, seipin could help recognize and stabilize pre-existing nascent droplets in the ER membrane (20, 21, 23). In both cases, however, the relationship between the role of seipin in LD formation and its ability to regulate lipid metabolism remains unclear.

Here, we use coarse-grain (CG) molecular dynamics (MD) simulations to investigate the mechanism of seipin in molecular detail. We find that seipin is able to cluster TG molecules inside its ring-like structure, and that both the transmembrane helices and the luminal domain contribute to this process. Diacylglycerol (DG), the lipid intermediate between TG and PA in the Kennedy pathway, also accumulates around seipin, further promoting the accumulation of TG at very low TG-to-phospholipids ratios. Our data suggest that by accumulating DG and TG molecules, seipin generates ER sites with a specific lipid composition, that in turn could promote the sequential recruitment of additional TG- and DG-sensing proteins involved in LD formation, including Promethin/LDOs, FIT/Yft2/Scs3 and perilipins.

## Results

### Seipin promotes the accumulation of TG molecules at low TG concentrations

To investigate the role of seipin in LD formation *in silico*, we embedded a model of the seipin 11-mer, consisting of its luminal structure and of its transmembrane (TM) helices (Figure 1A), in lipid bilayers composed of di-oleoyl-phosphatidylcholine (DOPC) with varying concentrations of TG molecules. Our model derives from the luminal structure of human seipin recently solved using cryo-EM(28) and our models for the protein and lipids are consistent with the coarse-grain (CG) SDK/SPICA force field (38–40).

**Figure 1.**
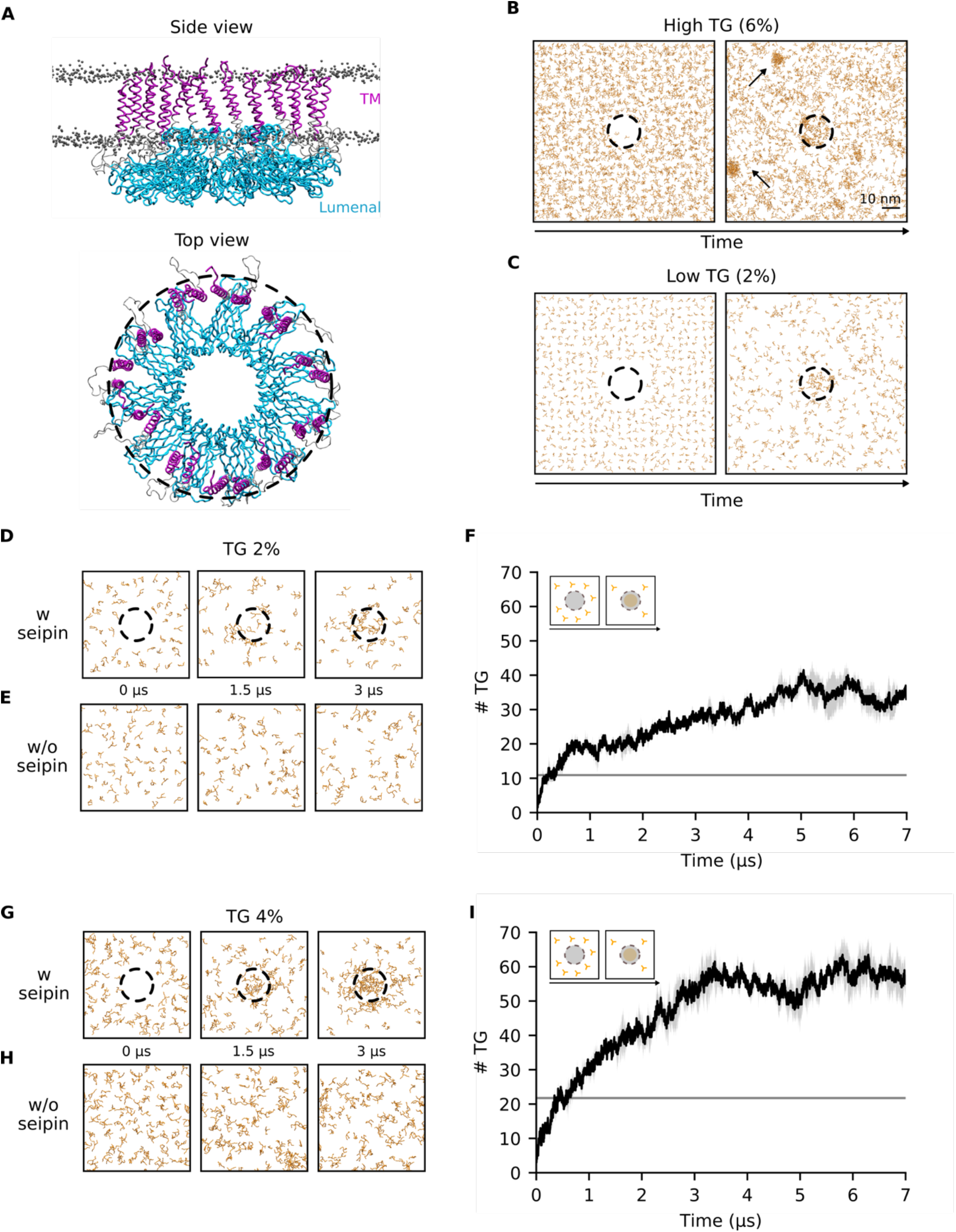
Seipin accumulates TG molecules in its ring-like structure at low TG concentrations. (A) Side and top view of the initial configuration of the MD simulations of the TM helices (purple) and luminal domain (cyan) of the human seipin undecamer embedded in a lipid bilayer. DOPC phosphate groups are shown as grey dots. (B-C) Top view of snapshots representing the time evolution of TG molecules in MD simulations at 6%(B) and 2%(C) TG concentrations. The black dotted ring represents the outermost radius of the seipin oligomer and indicates the presence of seipin in the simulations. TG molecules are colored in orange. (D,E) Top view of snapshots representing the time evolution of TG molecules in MD simulations at 2% TG concentration, with (D) or without (E) seipin. (F) Quantification of the number of TG molecules inside the seipin ring over time during the MD simulation depicted in (D). The grey line indicates the amount of TG molecules that would be present in an area of equivalent size of that measured for seipin if the TG molecules were distributed uniformly in the bilayer. (G,H) Top view of snapshots representing the time evolution of TG molecules in MD simulations at 4% TG concentration, with (G) or without (H) seipin. (I) Quantification of the number of TG molecules inside the seipin ring over time during the MD simulation depicted in (G). The grey line indicates the amount of TG molecules that would be present in an area of equivalent size of that measured for seipin if the TG molecules were distributed uniformly in the bilayer. Standard deviation of TG accumulation in different replicas (n=2) are shown as grey shading around the average value in panels E,G.

We first investigated whether seipin helps clustering TG molecules at low (2%) and high (6%) TG concentration. We chose these two concentrations as previous work(41, 42) showed that above 5% TG, spontaneous formation of TG blisters takes place in protein-free simulations, while this is not the case below this threshold. In agreement with those simulations, we observed that at higher concentrations (6% TG), spontaneous formation of TG blisters quickly (<300 ns) took place outside the seipin ring (Figure 1B). At low concentrations (2% TG), however, we invariably observed that TG molecules only accumulated in the proximity of the protein (Figure 1C).

To quantify the extent of seipin-induced TG accumulation at low TG concentration (2%), we computed the amount of TG molecules per seipin oligomer over time, and we compared it with protein-free simulations (Figure 1D-F). We observed that TGs accumulate in the presence of seipin (Figure 1D,F), reaching a plateau after few microseconds at approximately 35 TG molecules per seipin oligomer (Figure 1F). On comparable time scales, no such accumulation was observed in the absence of seipin (Figure 1E). To investigate whether the accumulation of TG molecules inside seipin is driven by specific protein-TG interactions or by demixing of TG molecules inside the lipid bilayer, we next estimated the amount of TG molecules inside the seipin ring that are in direct contact with the protein and compared it with those that do not interact with the protein, but rather are in contact with other TG molecules (Figure S1). Notably, 90% of the TG molecules inside the seipin ring establish direct contact with the protein, suggesting a protein-specific mechanism of TG accumulation.

To investigate whether the initial protein-bound TG molecules could act as seed for the growth of TG blisters via demixing of TG molecules from the bilayer, we repeated our simulations at a higher TG concentration (4%) (Figure 1G-I). At this concentration, spontaneous blister formation is not expected to occur in protein-free systems(41, 42), but the amount of TG molecules is substantially higher than that required for saturating seipin at 2% TG concentration according to Figure 1F. Indeed, also at this TG concentration we did not observe any spontaneous TG blister formation in the absence of seipin (Figure 1H), whereas TG accumulated inside the seipin ring in the presence of the protein (Figure 1G). Interestingly, the number of total TG molecules inside the seipin ring reached saturation at around 55 TG molecules (Figure 1I), a value significantly higher than that observed at lower (2%) TG concentration. Analysis of protein-TG contacts shows that while the amount of TG molecules in contact with the protein remains stable (Figure S1), the percentage of TG molecules inside the seipin ring that do not engage in interactions with the protein but rather form TG-TG interactions increases significantly (Figure S1).

Taken together, our results suggest that the seipin oligomer clusters TG molecules at low TG-to-phospholipids ratios. The finite size of the seipin oligomer sets a threshold on the amount of TG molecules that the protein can accumulate via direct protein-TG interactions, but those molecules can act as a seed for further growth of the TG blister inside the seipin ring. However, the observation that even in the presence of excess amount of TG, blister growth is arrested at relatively low amounts of TG molecules (at least in comparison with protein-free systems, where TG blisters can accumulate up to hundreds of TG molecules(42)) indicates that seipin is likely insufficient to promote extensive LD growth *per se*, in the absence of partner proteins(15, 17) or variations in membrane properties (43–45).

### The luminal domain of seipin is responsible for accumulating and retaining TG molecules

We next sought to investigate the molecular mechanism through which seipin accumulates TG molecules. To do so, we first investigated where TG molecules accumulate inside the seipin ring, and we specifically focused on the role of the individual components of seipin: the TM helices and the luminal region. These two domains are physically distinct in the seipin oligomer (Figure 2A,B), with the helices occupying the external ring of the complex and the luminal domain occupying the inner ring.

**Figure 2.**
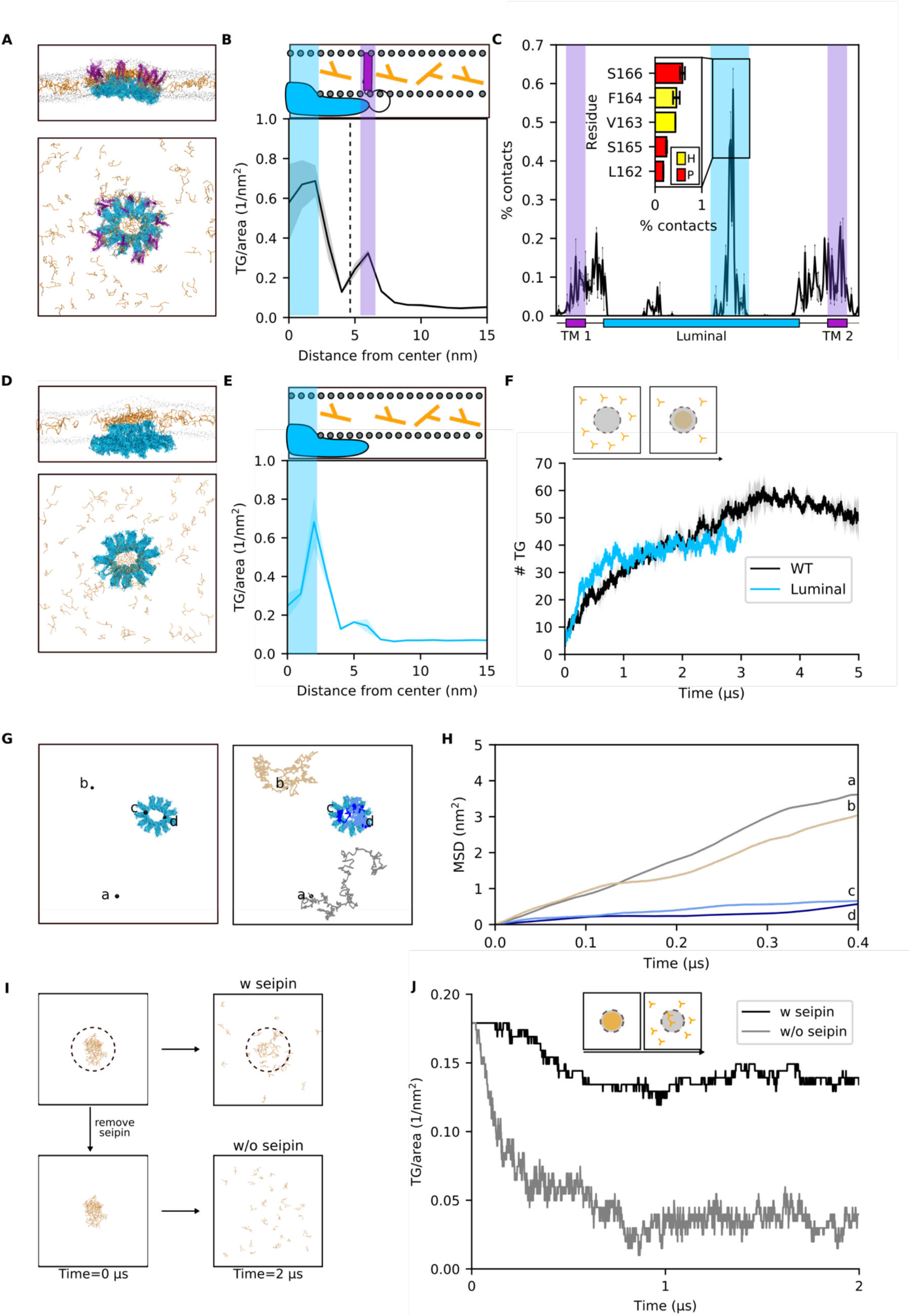
The luminal domain of seipin is sufficient to promote TG accumulation and retention. (A) Side and top view of the accumulation of TG molecules in the presence of seipin. The color codes are the same as in Figure 1. (B) Radial concentration of TG molecules in simulations of the entire seipin undecamer. (C) TG occupancy at seipin residues. The inset shows TG occupancy at the 5 protein residues that make the most contacts with TG molecules during the MD simulations. Color code: hydrophobic (yellow), polar (red). (D) Side and top view of the accumulation of TG molecules in the presence of the luminal domain of seipin alone. (E) Radial concentration of TG molecules in simulations of seipin luminal domain alone. (F) Quantification of the number of TG molecules inside the seipin ring over time during the MD simulation of seipin luminal domain. (G) Top view of the diffusion path of 4 TG molecules representative of the free diffusion in the lipid bilayer (brown and grey) and of the confined diffusion in proximity of the luminal domain of seipin (dark and light blue) (H) Corresponding mean square displacements. (I) Top view of initial and final snapshots of MD simulations of dissolution of the TG blister in the presence (top) or absence (bottom) of seipin. (J) Time evolution of the number of TG molecules inside the seipin ring when TG molecules are completely depleted from the surrounding lipid bilayer in the presence (black) or absence (grey) of seipin.

To do so, we first computed the distribution of TG molecules as a function of the radius from the center of the oligomer ring (Figure 2B). We observed the presence of two clear peaks at which TG molecules accumulate (Figure 2B). These two peaks appear at R ≈ 2.5 nm and R ≈ 6 nm and they represent, respectively, the extent of TG accumulation by the luminal and TM domains. Notably, TG accumulation is significantly higher in the proximity of the luminal domain in comparison with the TM helices (Figure 2B).

To further quantify the contribution of the two domains towards TG clustering, we next calculated the average residency time of TG molecules at specific protein interaction sites. To do so, we quantified the percentage of the trajectory during which the distance between a TG molecule and a specific residue was below 0.6 nm (Figure 2C). Consistently with our previous observation (Figure 2B), we found that TG molecules are in close contact with both TM helices and the luminal domain (Figure 2C). In addition, we could identify that the binding of TG molecules to the luminal domain takes place mainly via its HH, with residues V163, F164, S165 and S166 establishing the most contacts (Figure 2C, inset).

The analyses of Figure 2B,C indicate that the luminal domain is primarily responsible for the accumulation of TG molecules inside seipin. To investigate whether the luminal domain is sufficient to promote this process, we performed analogous simulations in the presence of the luminal domain alone (Figure 2D). Of note, in all simulations, the oligomeric structure of the protein was conserved during the simulations, a constraint which might not reflect physiological conditions in case some domains are also responsible for the oligomerization of the individual seipin monomers. We observed that in the absence of the TM helices, the luminal domain alone is able to promote TG accumulation comparably to that observed in the entire protein (Figure 2E, S2). Interestingly, the kinetics of TG accumulation is faster in the absence of the TM-helices (Figure 2F, S3).

Next, we investigated whether the TG molecules that are in contact with the luminal HH domain of seipin form stable contacts with the protein or rather exchange frequently with the surrounding lipid bilayer. To do so, we visually inspected the diffusion of TG molecules in contact with the protein and compared their diffusion with those that are free to diffuse in the bilayer (Figure 2G,H). Comparison of their mean square displacement clearly shows that molecules that engage in interactions with the luminal domain of seipin are restricted in their movement, and that they establish long-lived interactions with the protein (Figure 2H).

To further test the hypothesis of whether seipin, and specifically its luminal domain, is able to trap TG molecules in its proximity, we next investigated whether seipin is able to retain TG molecules when their concentration in the surrounding lipid bilayer is depleted. To do so, starting from the final state of our previous simulations in which TG molecules are clustered inside the seipin oligomer, we removed all free TG molecules that are not in direct contact with the luminal domain (Figure 2I). In these conditions, if seipin is also removed, TG molecules dissolve in the bilayer, and no cluster is present (Figure 2I,J). In the presence of the protein, on the other hand, only few TG molecules dissolve in the bilayer, while the vast majority remains stably associated to the protein (Figure 2J).

Taken together, our data suggest that the luminal domain of seipin, mostly through its HH region, is sufficient to accumulate TG molecules. The luminal domain is also responsible for the trapping of TG molecules inside the seipin ring, by establishing long-lived interactions that retain them. These molecules infrequently escape the protein even when the surrounding bilayer is depleted of TG. This suggests that seipin-positive ER sites might be enriched with TG molecules even in the absence of contemporaneous TG synthesis. Thus, while seipin diffuses in the ER membrane before LD formation(15), it might sequester TG molecules from the ER and drag them along in its diffusion path.

### The TM helices of seipin control the kinetics of TG accumulation and remodel the lipid composition at the periphery of the seipin ring

While the luminal domain of seipin appears to be sufficient to promote TG accumulation and to trap TG molecules (Figure 2), the time-averaged analyses of Figure 2B,C indicate that both the luminal domain and the TM helices contribute to TG clustering inside the seipin ring.

To better characterize the role of seipin TM helices in the process of TG accumulation by seipin, we first computed the time evolution of the radial concentration of TG molecules (Figure 3A). This analysis shows that TG molecules first accumulate around the TM helices, before proceeding towards the core of the ring and engaging in interactions with the luminal domain (Figure 3A). This suggests that the TM helices might play an important role in controlling the kinetics of TG accumulation.

**Figure 3.**
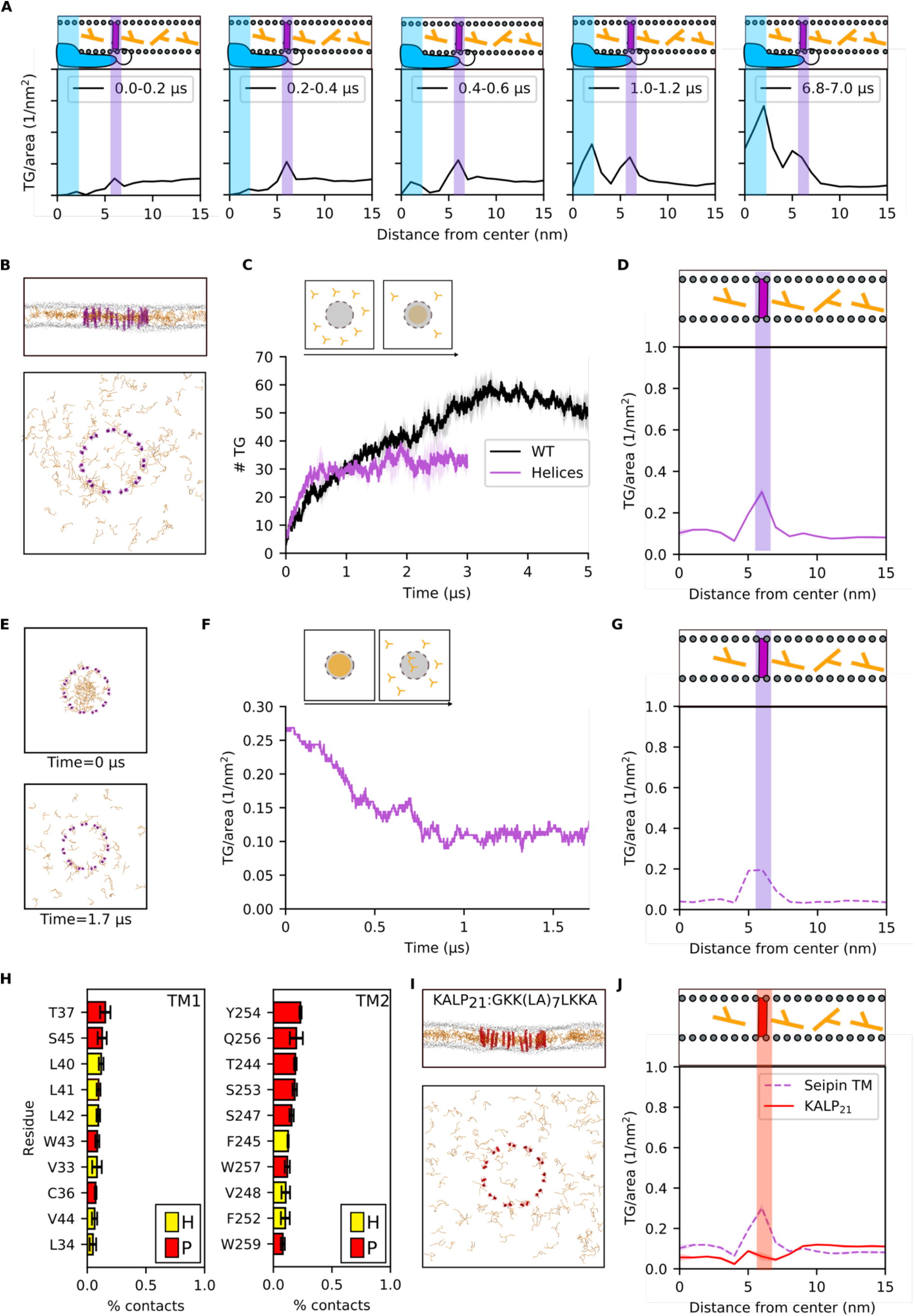
The TM helices of seipin control the kinetics of TG clustering and promote the accumulation of TG molecules at the periphery of the seipin ring. (A) Time evolution of the radial distribution of TG molecules inside seipin. (B) Side and top view of the accumulation of TG molecules in the presence of the TM helices of seipin alone (TM-only system). (C) Quantification of the number of TG molecules inside the seipin ring over time during MD simulation of the TM-only system. (D) Corresponding radial distribution of TG molecules (E) Top view of initial and final snapshots of MD simulations of dissolution of the TG blister in the TM-only system. (F) Time evolution of the number of TG molecules inside the seipin ring when TG molecules are completely depleted from the surrounding lipid bilayer in the TM-only system. (G) Radial distribution of TG molecules around seipin in MD simulations of blister dissolution in the TM-only system. (H) TG occupancy of the 10 protein residues in the TM helices that make the most contacts with TG molecules during the MD simulations. Color code: hydrophobic (yellow), polar (red). (I) Side and top view of the KALP_21_ system. (J) Radial distribution of TG molecules in the KALP_21_ system (red) in comparison with the TM-only system (purple).

To test this hypothesis, we investigated the process of TG accumulation in the presence of the TM helices alone (Figure 3B-D), and we compared its kinetics with that of the entire protein (Figure 3C). We observed that the initial kinetics is similar in both cases, before TG accumulation in the TM-only systems plateaus at a lower value in comparison with the entire protein (Figure 3C, S4). To further quantify this behavior, we computed the instances of TG molecules entering and exiting the seipin oligomer (Figure S5). We observed that the number of entry and exit events is comparable in the wild-type and in the TM-only system, but remarkably different in the luminal-only system, further suggesting that the TM helices, that are located in the external part of the seipin ring, effectively act as gates for the entry of TG molecules.

Next, we investigated whether the TM-only system is able to retain TG molecules in its center once a pre-existing TG blister is present (Figure 3E). Similarly to the protein-free system, the TM-only system fails to retain TG molecules within the blister (Figure 3F). Consistent with the observed higher frequency of TG entries and exits in the TM-only system, this lack of retention further indicates that the ring-like architecture alone is insufficient to prevent diffusion of TG molecules away from the protein (Figure S5). Analysis of the radial distribution of TG accumulation, however, shows that also in the simulation of blister dissolution (Figure 3E,F) in the TM-only systems, a small but non-negligible TG accumulation can be observed around the TM helices (Figure 3G). Of note, this TM-induced TG accumulation is identical with or without the luminal domain, as can be appreciated when comparing the peaks at 6 nm (Figure 3D,G and Figure 2B). Analysis of the residues making the most contacts between the TM helices and TG molecules, highlights the important role of multiple polar residues (T244, S247, S253, Y254, Q256,) that establish direct contact with the slightly polar glycerol moiety of the TG molecules (Figure 3H).

To further test whether the sequence of seipin TM helices is indeed specific for TG recruitment, we substituted the TM helices of seipin with artificial KALP_21_ peptides that lack polar residues in their sequence(46) (Figure 3I). In this condition, no accumulation of TG was observed (Figure 3J), further pointing to a specific role of polar residues in TG accumulation and indicating that the TM-induced TG accumulation at the periphery of the seipin oligomer might be important for its physiological role.

### Seipin is able to accumulate and trap diacylglycerols (DG), and DG enrichment doesn’t alter seipin’s ability to cluster TG molecules

Our simulations show that the seipin oligomer clusters and traps TG molecules, and that this activity is driven by its luminal domain. Even if the presence of the TM helices is not necessary towards this process, the TM helices of seipin distinctly promote local TG accumulation in their proximity by means of specific protein-TG interactions. We thus reasoned that this local membrane remodeling could be of relevance to the mechanism of seipin, including, possibly, to establish protein-protein interactions with other membrane-embedded proteins, especially as the TM helices represent the outermost portion of the protein oligomer.

We thus investigated whether lipids other than TG could accumulate around seipin. In particular, we focused on the precursors of TG along the Kennedy pathway, DG and phosphatidic acid (PA), as both have been shown to accumulate at LD formation sites in various physiological(31) or non-physiological conditions(29). Analysis of the local enrichment factor around seipin shows that DG molecules significantly accumulate inside the seipin ring (Figure 4A), while PA does not (Figure S6). Of note, DG accumulation is dramatically faster than that of TG, and many more DG molecules (up to 200) can be accommodated inside a single seipin oligomer (Figure 4B). Correspondingly, DG accumulation around both TM helices and luminal domain is significantly larger than that of TG (Figure 4C). Also, as is the case for TG molecules (Figure 2I,J), when excess DG molecules are depleted from the surrounding bilayer (Figure S7), DG molecules remains trapped inside the seipin ring, whereas they quickly dissolve in the bilayer if the protein is removed (Figure S7).

**Figure 4.**
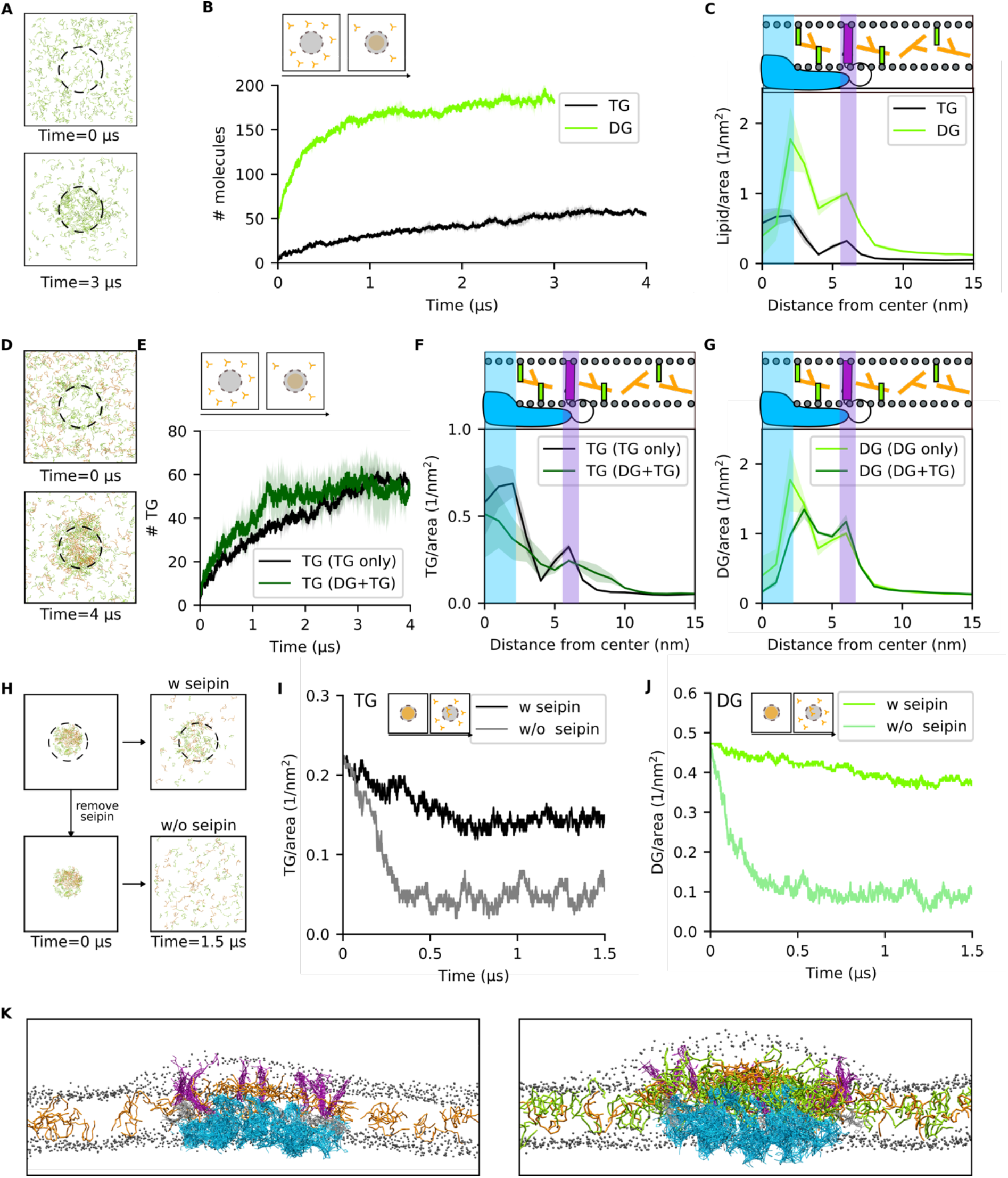
Seipin accumulates and traps diacylglycerols without losing its ability to cluster TG molecules. (A) Top view of snapshots representing the time evolution of DG (green) molecules in MD simulations in the presence of seipin. (B) Quantification of the number of DG molecules (green) inside the seipin ring over time in comparison with the TG-only (4% TG) system (C) Corresponding radial distribution of DG molecules in comparison with the TG-only (4% TG) system (D) Top view of snapshots representing the time evolution of DG (green) and TG (orange) molecules in MD simulations in the presence of seipin when TG molecules are randomly added to the lipid bilayer in the presence of pre-existing DG accumulation inside the seipin ring. (E) Quantification of the number of TG molecules inside the seipin ring over time when DG is already present in the system (dark green) in comparison with the TG-only system (black). (F) Corresponding radial distribution of TG molecules in the DG+TG (dark green) and in the TG-only system (black). (G) Corresponding radial distribution of DG molecules in the DG+TG (green) and in the DG-only system (light green). (H) Top view of initial and final snapshots of MD simulations of dissolution of the TG+DG blister with (top) and without (bottom) seipin. (I,J) Time evolution of the number of TG (I) and DG (J) molecules inside the seipin ring when TG and DG molecules are completely depleted from the surrounding lipid bilayer in the presence (black, green) or absence (grey, light green) of seipin (K) Side view of snapshots representing the equilibrated conformation of in-bilayer seipin in the presence of TG-only (left) or DG+TG (right). Color code: seipin luminal domain (blue), seipin TM domains (purple), TG (orange), DG (green), DOPC phosphate group (grey).

As DG is the precursor of TG, we next investigated whether the TG accumulation observed in the absence of DG is promoted or abolished by the pre-clustering of DG molecules inside the oligomer ring (Figure 4D). We observed that in the presence of DG, TG molecules still accumulate inside seipin (Figure 4D), and they do so with comparable kinetics, if not faster (Figure 4E). Remarkably, the amount of TG molecules that accumulate inside the seipin ring is analogous to those observed in DG-free simulations (Figure 4E), despite the initial presence of DG molecules in contact with the protein. Of note, however, when both DG and TG molecules are present in the system, TGs are less likely to interact directly with the protein (Figure 4F, S8), but rather distribute somewhat uniformly inside the seipin ring, with a maximal concentration in the center of the ring (Figure 4F). In parallel, the presence of TG molecules slightly displaces DG molecules from the luminal domain of seipin, further increasing DG concentration close to the TM helices (Figure 4G).

Finally, we tested whether seipin is able to retain both DG and TG molecules when they are depleted from the surrounding membrane (Figure 4H). Also in this case, both DG and TG molecules remain trapped inside the protein ring (Figure 4I,J).

Taken together, our data suggest that the seipin oligomer is able to significantly cluster DG molecules, and that the presence of DG further promotes the ability of seipin to cluster TGs. The simultaneous presence of both DG and TG molecules in the seipin ring leads to a dramatic enhancement in overall blister size with respect to the presence of only TG molecules (Figure 4K). As a consequence, the presence of seipin together with the associated TG and DG blister substantially remodel the surrounding bilayer in two distinct ways: first, it promotes a significant remodeling of the lipid composition in its surroundings, with a marked DG accumulation in its vicinity; second, it induces a non-negligible positive curvature to the cytosolic leaflet of the lipid bilayer (Figure 4K, S9).

## Discussion

Identified as a crucial protein in LD biogenesis and homeostasis many years ago(47), the molecular mechanism of seipin has remained enigmatic and debated(14–17, 20–22, 24, 28, 29, 34, 48). Here, we took advantage of the recently solved cryo-EM structures of the luminal domain of seipin (28, 34) to perform MD simulations of the seipin oligomer embedded in explicit lipid bilayers enriched in TG.

Our simulations suggest that seipin can cluster TG molecules even when they are present at low concentration in the surrounding membrane, and that it does so by first binding to them through several polar residues located in its TM helices. These interactions are short-lived, and TG molecules can subsequently diffuse towards the interior of the ring and engage in stable interactions with the HH of the luminal domain, from which they hardly detach. These strongly-bound TG molecules can act as nucleation seeds for newly arriving TG molecules, further promoting TG accumulation. In addition, we found that a similar mechanism is true for the TG precursor, DG, and that accumulation of DG inside seipin does not prevent the subsequent accumulation of TG.

While these observations suggest a mechanistic role for seipin in the early stages of LD formation, the presence of seipin alone seems to be insufficient to promote TG accumulation beyond a certain size threshold, as in our simulations TG blister growth reaches a plateau even in the presence of excess TG molecules in the lipid bilayer. This observation could also explain why TG was not identified to co-precipitate with seipin in the absence of its partner protein LDAF1(15), given that a limited accumulation of TG molecules paired to minor loss during the purification process could produce a signal below the detection limit.

Rather, our simulations suggest that the membrane remodeling activity of seipin might play a major role in the subsequent recruitment of additional proteins that are required for the completion of LD biogenesis(15–19) and its regulation(11, 33, 49). To this extent, the unexpected observation that seipin also accumulates DG, and that it does so even more efficiently than TG, is extremely intriguing. Unlike TG, DG is a constitutive ER lipid that is generally present at non negligible concentrations and that it has been shown to promote the recruitment of several LD-associated proteins, including perilipins(50), FIT proteins(51), and the TG-synthesizing acyltransferase Lro1(31). Notably, our results are in agreement with the recent observation that seipin-positive ER sites are also enriched in DG, even in the absence of LD formation (31), and they suggest that seipin could be directly responsible for the observed DG enrichment via direct accumulation of DG molecules in its structure. Also, our observation that seipin, upon TG and DG accumulation, induces a positive curvature to the lipid bilayer, and specifically to the cytosolic leaflet, not only agrees with the experimental observation that seipin preferentially localizes in curved regions of the ER (52), but also suggests that the generation of local curvature might help in both the recruiting of downstream protein partners (e.g. LDAF1, perilipins…) and in providing the correct directionality for subsequent LD growth and budding.

In addition to its role in LD biogenesis, our simulations hint at an intriguing possible role of seipin in maintaining ER homeostasis: as it diffuses through the ER bilayer in its oligomeric assembly(15), seipin would be able to remove potentially lipotoxic TG (and DG) molecules from the ER by accumulating them in its structure, where they could subsequently promote protein recruitment and act as a seed for the biogenesis of LDs. We expect that future studies will address the role of seipin in ER homeostasis, and whether the seipin mechanism we have identified is specific to DG and TG accumulation, or if it is also valid for other neutral lipids and their precursors, such as steryl-ester and cholesterol.

Finally, we want to highlight the limitations of our modeling approach. First, our seipin model lacks the N-terminal domain (residues 1-21). We opted not to model this domain *ab-initio* as it has been shown that while deletion of the first 14 amino acids at the N terminus of seipin results in a delay in LD formation, this doesn’t affect LD morphology(24). In addition, in our model, only the luminal part of seipin was built using experimentally-derived coordinates, while the TM-helices were modeled *ab initio* as straight helices based on the coordinates of the cryo-EM structure of the luminal domains. While this leaves room for improving the accuracy of the TM helix region model with respect to experiment, the observation that the luminal domain alone is sufficient for the accumulation and trapping of TG molecules indicates that the bias introduced in any potentially erroneous modeling of the TM helices is likely insufficient to alter the main conclusions of our work. Second, the use of CG modelling introduces inherent quantitative shortcomings in the accurate reproduction of molecular interactions, most notably in the case of charged or polar residues where electrostatic effects or hydrogen bonding might play a major role. In our case, this might be particularly important pertaining to the observation that seipin interacts with TG molecules via multiple polar residues. However, a specific role of the polar moiety of TG in establishing interactions with proteins is supported by recent experimental observations(53) and could be the subject of future validation by atomistic simulations. Overall our simulations provide a high-resolution description of the mechanism through which seipin promotes LD formation, and they indicate that seipin ER remodeling activity is a crucial component of its molecular mode of action. We foresee that future studies including additional proteins that are known to interact with seipin at sites of LD biogenesis will shed further light into the entire process of LD formation and growth with unprecedented molecular detail.

## Author Contributions

SV and VZ designed the study with the help of WS, VZ performed MD simulations. WS contributed the protein force-field, VZ, and SV analyzed the computational data. VZ and SV wrote the manuscript with the help of WS.

## Acknowledgments

This work was supported by the Swiss National Science Foundation (grant #163966). This work was supported by grants from the Swiss National Supercomputing Centre (CSCS) under project ID s726 and s842 and s980. We acknowledge PRACE for awarding us access to Piz Daint, ETH Zurich/CSCS, Switzerland. VZ acknowledges the JSPS/ETH young researchers’ exchange program (Fellowship # GR19107).

## Methods

### Molecular dynamics (MD) simulations

All MD simulations were performed using the software LAMMPS (Plimpton, 1995) in combination with the Shinoda-Devane-Klein (SDK)(39, 40, 54–56) and the SPICA force field(57) for proteins (manuscript in preparation) based on the previous parameterization of the individual amino acids(58). Combination rules were used to derive non-bonded interactions of TG and DG with the protein. Initial configurations and input files were obtained through conversion of atomistic snapshots using the setup lammps (https://www.spica-ff.org/index.html) or CG-it (https://github.com/CG-it/) tools and in-house scripts.

In the simulations, temperature and pressure were controlled via a Nosé-Hoover thermostat(59) and barostat (60–62). Target temperature was 310K and average pressure was 1 atm. The lateral xy dimensions were coupled, while the z dimension was allowed to fluctuate independently. Temperature was dumped every 0.5 ps, while pressure every 5 ps. Linear momentum was removed every 1 timesteps. Van der Waals and electrostatic interactions were truncated at 1.5 nm. Long-range electrostatics beyond this cutoff were computed using the particle-particle-particle-mesh (PPPM) solver, with an RMS force error of 10^−5^ kcal mol^−1^ Å^−1^, order 3 and a grid size of 16, 16, and 18 Å in the x, y, and z directions, respectively. A time step of 10 fs was used, except for simulations without the protein in DOPC/TG bilayers, where a timestep of 20 fs was used. An initial equilibration was carried out by performing energy minimization, followed by a short NVT run of 100 ps and a short NPT run of 100 ps. In the simulations containing only helices (both seipin and KALP_21_), positional restraints along the x and y direction were used in order to preserve the ring structure, with a force constant of 10 kcal mol^−1^ Å^−2^.

### MD systems setup

To build the model of the protein used in this work, the structure of the luminal portion of the human seipin (ID:6DS5)(28) was taken from RCSB PDB. Furthermore, our model includes the transmembrane (TM) domains of the protein (two per monomer), and the loops connecting the TM domains with the luminal domains. To build these domains, we first modelled the TM domains (residues 27-47 and 243-263) as fully folded alpha-helices using VMD(63). We next positioned the TM helices above the last residues solved in the crystal structure using Packmol(64). From this configuration we modelled the loops connecting the TM domains and the luminal domains using the MODELLER tool ModLoop(65). We also added 5-6 residues (residues 21-26, 263-269) on the N- and C-terminus of the protein to improve the overall stability of the model. Overall, our final model consists of the undecameric oligomer comprising the TM helices, the luminal portion and the loops connecting them (residues from 21 to 269). This structure was embedded in a dioleoyl–phosphatidylcholine (DOPC) bilayer using CHARMM-GUI(23) and equilibrated with 10 ns all-atom MD simulations. The final coordinates were converted to CG and used as starting point for the CG simulations. In systems containing only the luminal part of the protein, the cryo-EM structure was used as a starting point.

In systems containing only the seipin TM helices, the same approach described above was used, but the positioned helices were not connected to the rest of the protein. A similar approach was used for the KALP_21_ (sequence GKK(LA)_7_LKKA) mutant, where 22 helices were modeled and positioned in the same position as the WT helices. In all cases, the proteins were converted to CG and an elastic network with a cutoff of 0.9 nm for the intramolecular elastic bond and a force constant of 1.195 kcal mol^−1^ Å^−2^ was used to keep the secondary structure fixed.

In order to study the dissolution of TG blisters upon protein removal, we selected the coordinates of equilibrated simulations containing seipin and we subsequently removed all the molecules outside a radius of 8 nm from the center of the protein. In the systems “without seipin”, the protein was also removed. All the systems were neutralized and ionized to a concentration of 0.15 mol/L using the GROMACS(66) tool genion. Multiple independent replicas were run and details of the systems are reported in Table 1.

**Table 1.** Details of the molecular simulations presented in this work. The abbreviation “WT” denotes the presence of the entire seipin protein in the system, “L” of only the luminal part and “H” of only the TM region.

| *Accumulation* |  |  |  |
| --- | --- | --- | --- |
| Bilayer composition<br>(# of molecules) | TG<br>molecules | # of<br>replicas | Length<br>(μs) |
| 3200 DOPC | 64 | 1 | 7 |
| 3200 DOPC | 128 | 1 | 7 |
| 28773 DOPC + S (big) | 545 | 1 | 2.5 |
| 28492 DOPC + S (big) | 1704 | 1 | 0.3 |
| 3281 DOPC + WT | 110 | 2 | 7 |
| 3281 DOPC + WT | 57 | 2 | 7 |
| 3368 DOPC + L | 110 | 2 | 3 |
| 3357 DOPC + H | 115 | 2 | 3 |
| 3357 DOPC + H (KALP) | 115 | 2 | 3 |
| 2576 DOPC + 310 DOG + WT | 0 | 2 | 3 |
| 2576 DOPC + 310 DOG +WT | 110 | 2 | 4 |
| 2911 DOPC + 344 POPA +WT | 0 | 2 | 2.5 |
| *Dissolution* |  |  |  |
| 2576 DOPC + 95 DOG + WT | 45 | 1 | 1.5 |
| 2576 DOPC + 95 DOG | 45 | 1 | 1.5 |
| 2756 DOPC+125 DOG + WT | 0 | 1 | 1 |
| 2756 DOPC+125 DOG | 0 | 1 | 1 |
| 3661 DOPC +WT | 36 | 1 | 2 |
| 3661 DOPC | 36 | 1 | 2 |
| 2771 DOPC + H | 54 | 1 | 1.7 |

### Simulations analysis

In the systems containing seipin, the accumulation of TG molecules as a function of distance from the center of the seipin ring was calculated by counting the number of lipid molecules in annuli delimited by concentric circles of radius=R+1 nm.

The accumulation of DG/TG/POPA (lipid) molecules over time in seipin proximity was computed by summing the number of lipid molecules in annuli with a radius=R+8 nm. Lipid/area calculation was obtained by dividing the total number of lipid molecules by the total area of the annulus. The same calculation was used to calculate the number of molecules dissolved in the bilayer in the setup “dissolution”, but counting only the lipid molecules within a radius of 8 nm from the center of the seipin ring, with the exception of Figure S3 and S4 for which a radius of 6 nm was used.

For the analysis of the contacts between the protein and TG molecules, the number of contacts between each residue of the protein and the glycerol bead of TG were calculated using the software GROMACS(66). A contact was defined when the distance between TG and the protein is lower than 0.6 nm. The number of contacts for corresponding residues in different monomers was summed and the final values are normalized by the length of the simulations.

The number of contacts between the protein and TG was calculated with the same criterium described above. The percentage of TG-TG contacts interacting inside the ring of seipin was obtained by subtracting the numbers of protein-TG contacts to the number of TG molecules inside the protein (radius of distance from the seipin center <=8 nm). Only the contacts for the last 2 μs of simulation are averaged.

Trajectories of single molecules for Figure 2G were obtained extracting the coordinates of TG molecules via the tool MDtraj(67) and used to plot the time evolution of the trajectory of the molecules and to compute the mean squared displacement.

For the calculation of the events in/out of the seipin ring, the coordinates of TG molecules from 3 μs of trajectory from different systems were extracted using the tool MDtraj and processed to count events. A buffer region with a radius between 6 and 8 nm from the seipin center was defined and an “in”-event was considered only if a TG molecule moved from a region of radius > 8 nm to a region of radius < 6 nm from the seipin center, or vice-versa for “out”-events.

To visualize the effect of different bilayer compositions and of seipin on bilayer curvature, we first aligned all systems so that the center of mass of the phospholipids is at {0,0,0}, and we then represented the head group of the phospholipids based on their z position (red negative values, white = 0, blue positive values).

All results are represented as an average and standard deviation of the two independent replicas, except for the systems “dissolution” for which only one replica was run.

All the images are rendered using VMD(63). The calculation of the number of TG molecules per distance from the protein seipin were performed using a tcl script, while the rest of analysis and graphs were produced using python scripts.

## Supplementary information

**Figure S1.**
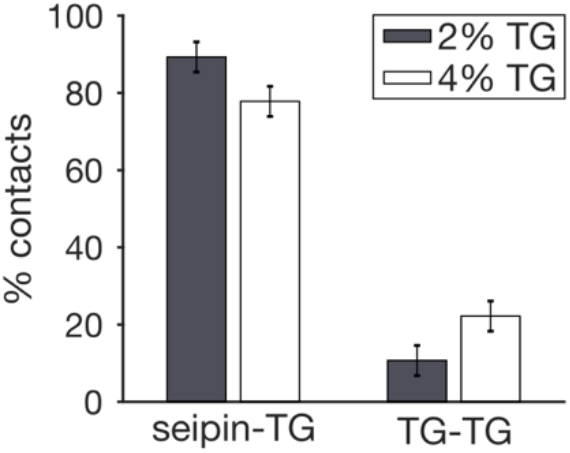
Percentage of seipin-TG or TG-TG in simulations at different TG concentrations.

**Figure S2.**
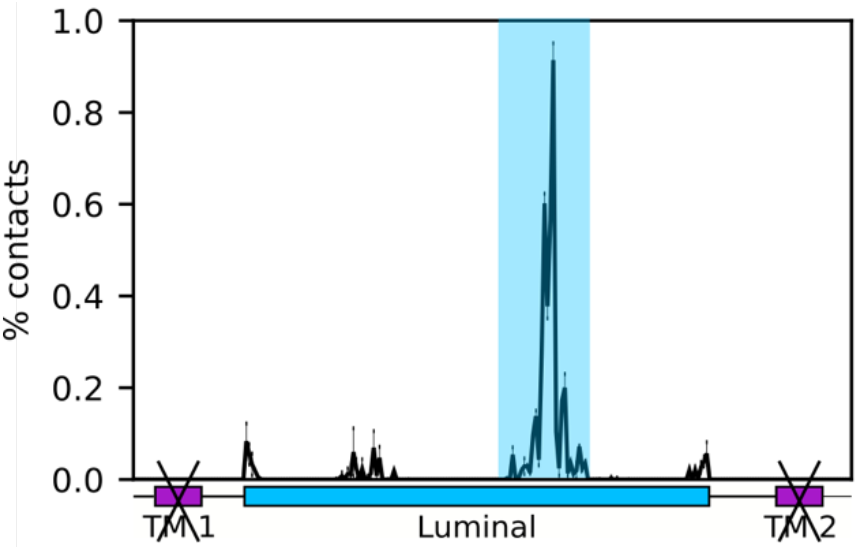
Percentage of contacts between the luminal part of seipin and TG in simulations containing only the luminal domain of seipin.

**Figure S3.**
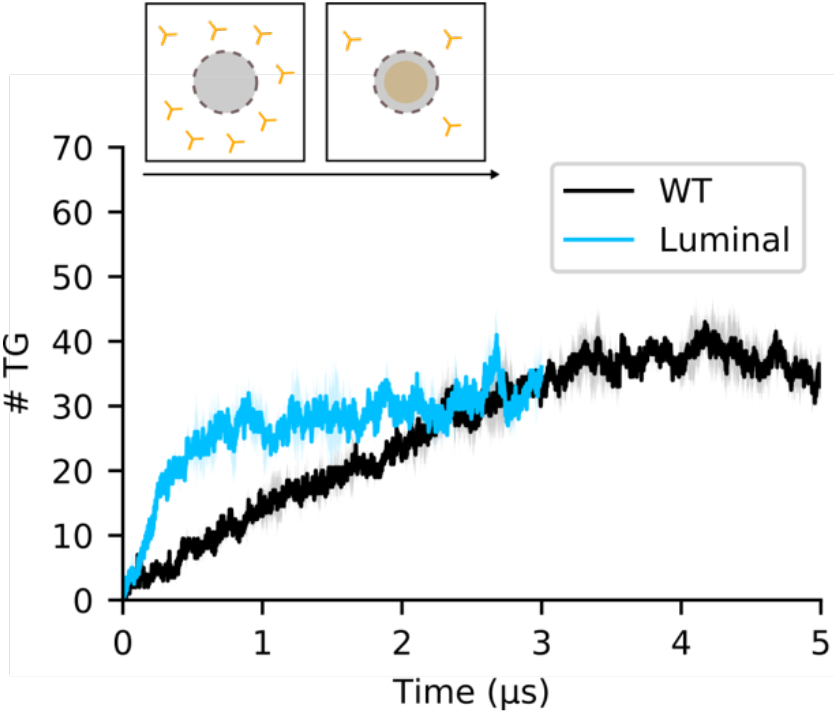
Number of TG molecules inside the inner part of the seipin ring (R= 6 nm) over time during the MD simulation of the luminal domain of seipin.

**Figure S4.**
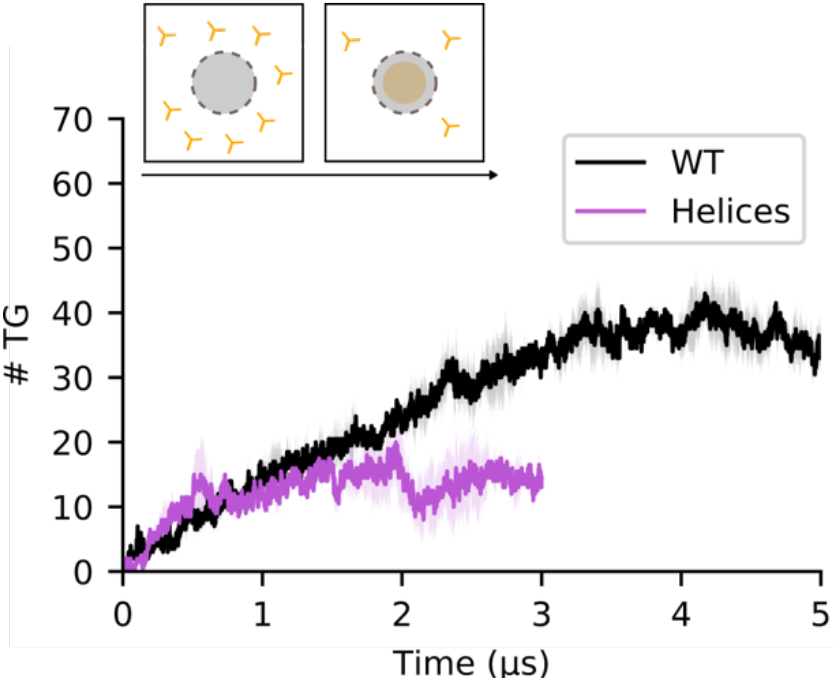
Number of TG molecules inside the inner part of the seipin ring (R= 6 nm) over time during the MD simulation of the TM domain of seipin.

**Figure S5.**
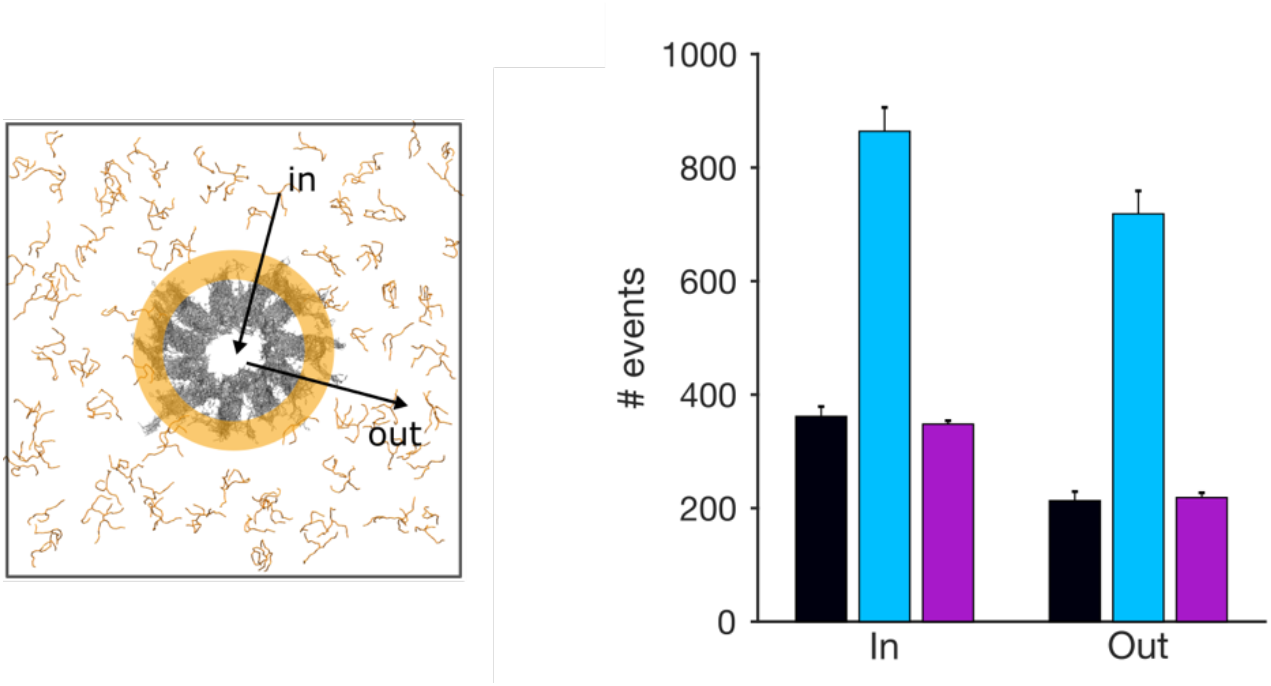
Number of events of molecules entering inside (in) or going outside (out) the seipin ring. The buffer region is colored in orange (see Methods section).

**Figure S6.**
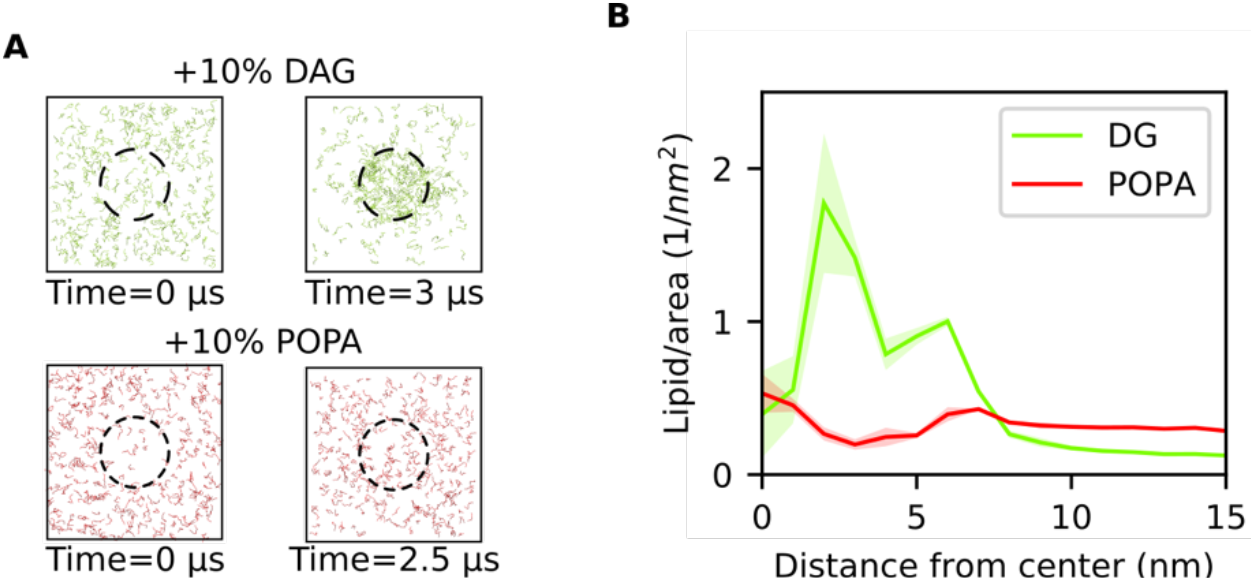
(A) Top view of systems containing DG or POPA at the beginning and at the end of the simulations. (B) Radial concentration of TG molecules in simulations of seipin with DG (green) or POPA (red).

**Figure S7.**
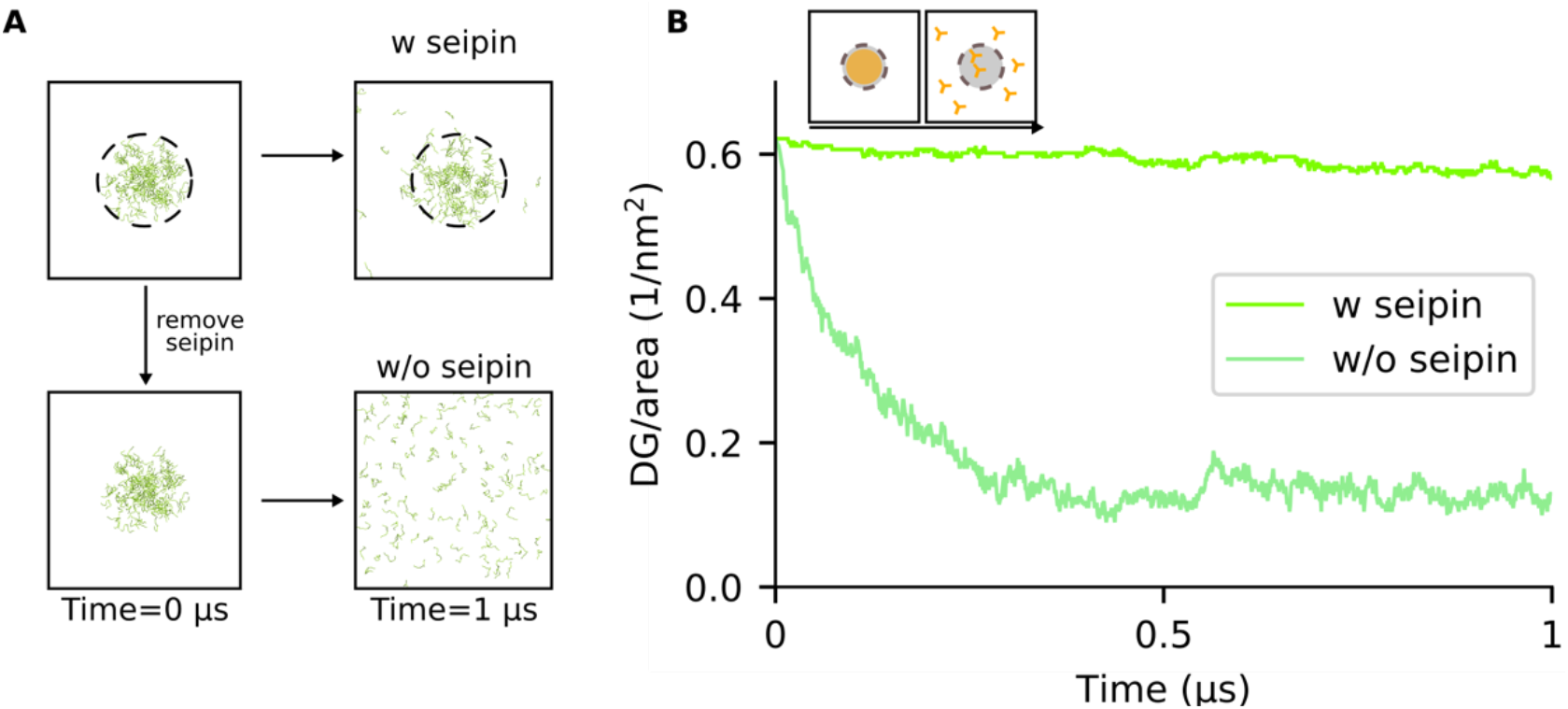
(A) Top view of initial and final snapshots of MD simulations of dissolution of DG blister with (top) and without (bottom) seipin. (B) Time evolution of the number of TG molecules inside the seipin ring when DG molecules are completely depleted from the surrounding lipid bilayer in the presence of the TM helices of seipin.

**Figure S8.**
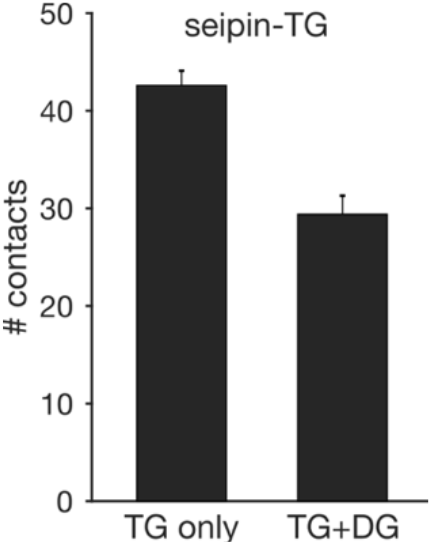
Number of contacts between seipin and TG in simulations containing only TG or in presence of DG.

**Figure S9.**
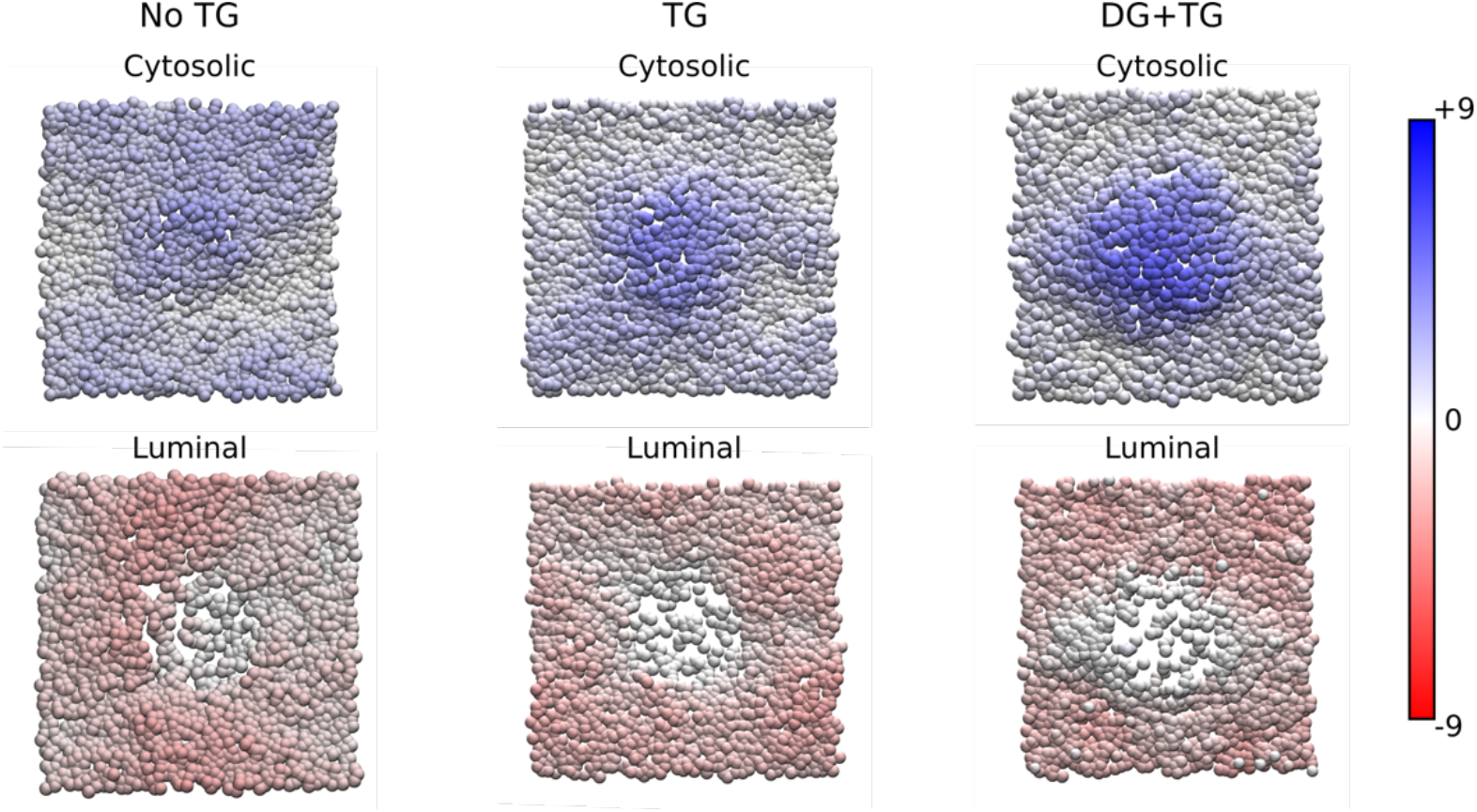
Top views of equilibrated snapshots of cytosolic and luminal leaflet in different systems (no TG, TG, DG+TG). The colors represent the position of the DOPC headgroups along the z direction (perpendicular to the membrane plane). Distances in the color scale are expressed in nm.

